# Two-color super-resolution localization microscopy via joint encoding of emitter location and color

**DOI:** 10.1101/2021.10.03.462920

**Authors:** Yujie Wang, Weibing Kuang, Mingtao Shang, Zhen-Li Huang

## Abstract

Multi-color super-resolution localization microscopy (SRLM) provides great opportunities for studying the structural and functional details of biological samples. However, current multi-color SRLM methods either suffer from medium to high crosstalk, or require a dedicated optical system and a complicated image analysis procedure. To address these problems, here we propose a completely different method to realize multi-color SRLM. This method is built upon a customized RGBW camera with a repeated pattern of filtered (Red, Green, Blue and Near-infrared) and unfiltered (White) pixels. With a new insight that RGBW camera is advantageous for color recognition instead of color reproduction, we developed a joint encoding scheme of emitter location and color. By combing this RGBW camera with the joint encoding scheme and a simple optical set-up, we demonstrated two-color SRLM with ∼20 nm resolution and < 2% crosstalk (which is comparable to the best reported values). This study significantly reduces the complexity of two-color SRLM (and potentially multi-color SRLM), and thus offers good opportunities for general biomedical research laboratories to use multi-color SRLM, which is currently mastered only by well-trained researchers.

## 1. Introduction

Multi-color super-resolution localization microscopy (SRLM) brings rich insights into the spatial relations and fundamental interactions among subcellular structures, which are beneficial for solving important questions in life sciences [1-3]. Generally, multi-color SRLM requires to label biological targets using emitters with distinct emission colors, where the ultimate goal is to distinguish these emitters with minimal crosstalk [4]. The reported methods for achieving this goal rely on different strategies, including mainly sequential excitation [5, 6], spectral splitting [7, 8], chromatic dispersion [4, 9], and point spread function (PSF) engineering [10, 11]. Sequential excitation methods sequentially capture different emitters, and distinguish these emitters from temporal image series. Spectral splitting methods capture different emitters simultaneously, use filters to split their emissions into several pathways, and finally recognize the emission colors using images from different cameras or different regions of the same camera. These methods usually suffer from medium to high crosstalk among different types of emitters, and thus can only distinguish emitters with large spectral separation [4-8].

Alternatively, fluorescence emission can also be dispersed into a set of adjacent pixels using a diffraction grating or prism [4, 9], and the color of the emitter is determined later through the degree of dispersion. This chromatic dispersion based multi-color SRLM method is suitable for distinguishing emitters with low crosstalk, even for those emitters with close spectral separation. However, this method usually requires a dedicated optical system (consisting of dichroic mirror, mirrors, fluorescence filters, and even multiple objectives) [6, 12] and a complicated image analysis procedure [4, 12], because image registration is required to align different types of emitters. On the other hand, PSF engineering [10, 11] includes emitter color into PSF with a phase modulation device, and requires only one detection pathway. This one-pathway method simplifies the optical set-up and image analysis procedure, but can only distinguish spectrally well-separated emitters with relatively high crosstalk.

Considering the limitations of current methods, we tried to find a simple and efficient method to realize multi-color SRLM. In this method, we aimed to reduce significantly the complexity in optical setup and image analysis, but still keep the desirable performance of low crosstalk to a level that is comparable to the best reported values [4, 5, 8]. In this way, we could promote more biomedical research laboratories to focus on challenging questions (e.g. particle tracking [13], relative distribution of cellular structures [14], and molecular counting [15]) that can be best investigated using multi-color SRLM. Fortunately, we noticed that a special type of scientific Complementary Metal-Oxide-semiconductor (sCMOS) camera, called RGBW camera [16, 17], may be used to achieve this goal. Note that RGBW camera typically contains filtered (or color) pixels and unfiltered (or white, W) pixels, and that color pixels includes mainly red (R), green (G), blue (B), and near infrared (NIR) pixels. RGBW camera was originally designed to perform color reproduction for low light photography, but is not successful in the consumer market. However, instead of color reproduction, we may use the color pixels to recognize emitter colors, and the white pixels to identify emitter location (for SRLM). Since the emission from an emitter covers a good number of color pixels (depending on the PSF size, typically >10), the emitter color may be well captured and later recognized from dissecting the relationships among these color pixels, thus enabling low crosstalk in classifying different types of emitters. That is to say, exploring the potentials of RGBW camera in color recognition rather than color reproduction may bring new possibilities in various application fields of multi-color SRLM.

Here we present a simple method for two-color (and potentially multi-color) SRLM via joint encoding of emitter location and color. To the best of our knowledge, the optical setup of our method is the simplest as compared to the reported two-color SRLM methods [4-11]. We designed a pixel-level scheme for joint encoding emitter location and color by taking advantage of the repeated pixel pattern (containing R, G, B, NIR, and W pixels) in a customized RGBW camera. Notably, this kind of customized camera is different from traditional monochrome (black-and-white) and color cameras, and the color pixels are used here for color recognition rather than color representation. Therefore, we prefer to rename it to colorimetry camera. Using this camera, we are able to encode both emitter location and color into a single raw image. Furthermore, by replacing a monochrome camera (which is traditionally used in multi-color SRLM) with this colorimetry camera and choosing multiple emitters which could be recognized by this camera, we would possibly perform multi-color SRLM through a simple optical set-up. As a pilot study, here we demonstrated the power and usage of this colorimetry camera in two-color SRLM with low crosstalk. Imaging with more colors is possible if we select more types of emitters, since the color pixels response to a broad spectral range.

## 2. Material and Methods

### 2.1 Sample preparation

Cos-7 cells were cultivated on 35-mm glass-bottom dishes. After overnight growth, samples were washed with PBS at room temperature and soaked with fixation buffer (3% paraformaldehyde, 0.05% glutaraldehyde and 0.2% Triton X-100 in PBS) for 15 minutes. After washed three times with PBS, cells were permeabilized and soaked with blocking buffer (3% BSA and 0.2% Triton X-100 in PBS) for 30 min. Cells were further stained with primary and secondary antibody successively at room temperature for 1 hour, washed three times with blocking buffer, washed with PBS, and then stored at 4 °C for further use. The primary antibodies were mouse monoclonal anti–α-tubulin antibody (T5168, Sigma), or/and rabbit anti-Tom20 antibody (HPA011562, Sigma). The secondary antibodies were DL633 labeled Goat anti-Mouse IgG (A-21235, Invitrogen) or/and CF680 labeled donkey anti-rabbit IgG (20820, Biotium).

### 2.2 Optical setup and imaging

Cell samples were soaked in standard SRLM buffer [18], and then imaged on a home-built SRLM system (Fig. 1(a)) based on an Olympus IX73 microscope. Cells were excited by a 640 nm laser (3W, LWRL640, Laserwave, China). Fluorescence emission was collected by a 60X/NA1.42 oil immersion objective (Olympus), transmitted through a dichroic mirror (ZT405/488/532/640rpc-XT, Chroma), focused by the tube lens, and then filtered with a band-pass filter (ET705/100m, Chroma). The filtered emission was finally captured by a customized RGBW camera (Retina 200DSC, Tucsen Photonics. Pixel size: 6.5 μm. Read noise: 2.71 e-rms) with an exposure time of 30 ms. The emission from different emitters (Fig. 1(d)) was imaged through the optical system in Fig. 1(a), captured by the color pixels (R, G, B, NIR pixels) and unfiltered pixels (W pixels) in the camera. The repeated pixel pattern of the camera is shown in Fig. 1(b). Emitter locations (see the dots in Fig. 1(d), and the crosses in Fig. 1(e)) can be encoded with W pixels, using PSF of the emitters. Emitter color can also be encoded into the same raw image (Fig. 1(e)), using the sensitivity changes in the color pixels under different wavelengths (Fig. 1(c)).

**Fig. 1.**
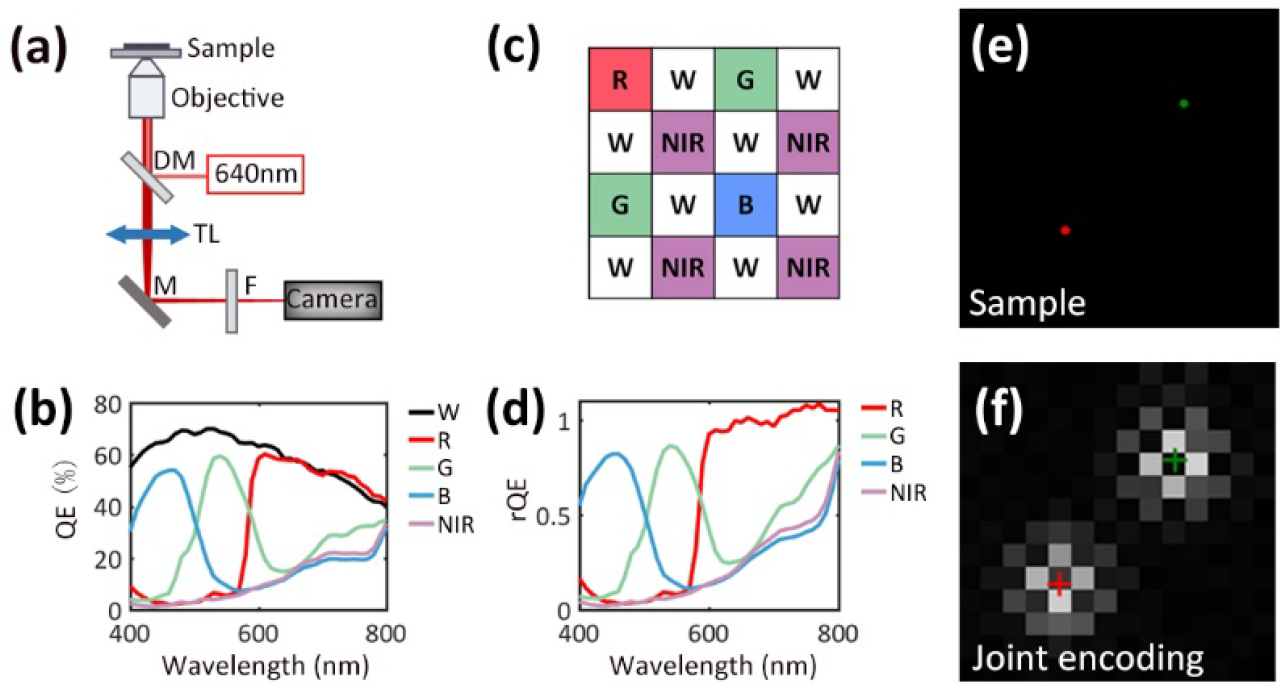
Schematic of multi-color SRLM via joint encoding of emitter location and color. (a) Optical set-up. DM: dichroic mirror; TL: tube lens; M: mirror; F: filter. (b) The spectral sensitivity curves of the W channel and the color channels in the colorimetry camera. Note that at some wavelengths, the QE values in the R channel are higher than those in the W channel. This is possibly due to different levels of transmission attenuation from the filling materials between the on-chip lens and the wiring. Color recognition in this study is based on experimentally measured NCI distributions, rather than these QE curves. (c) The repeated pixel pattern of the colorimetry camera, including four color channels (R, G, B, and NIR) and a W channel. (d) The sensitivity changes (or relative quantum efficiency, rQE) of the color pixels under different wavelengths, where rQE is the QE ratio between color channel and W channel: rQE= QE_color_/QE_W_. (d) Illustration of a sample consisted of two emitters with different color. (e) Illustration of an acquired image with joint encoding of emitter locations and color. The crosses indicate the original true positions of the emitters.

### 2.3 Determining emitter location for SRLM

For sparse emitters in SRLM, we calculate emitter locations from joint encoded raw images using two steps: subregion extraction and localization.

#### Subregion extraction

In a joint encoded raw image, the emission from an emitter is distributed in a subregion of pixels, where each pixel is associated with a channel (R, G, B, NIR, or W). The channel arrangement can be found in the repeated pixel pattern of the camera. Among these channels, we use only the W pixels to perform localization analysis. Hence, we design the following mask to extract W pixels from a raw image:

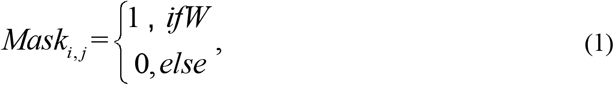

where *i, j*∈ [−*R, R*], and 2R+1 is equal to the subregion size.

To minimize the influences of noises and uneven background, we smooth the raw image with a Gaussian filter combined with the mask shown above. The filter has the following kernel:

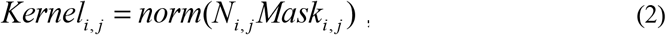

where N_i,j_ is Gaussian function, and norm means normalization of the sum in the subregion to 1. The standard deviation of the Gaussian function depends on the PSF of the optical system, and is set to be 1.3 pixels in this study.

A pixel can be detected as the center pixel of an emitter, if the intensity of this pixel is larger than the predetermined threshold and this pixel has the maximum intensity in the subregion. Pixels detected as center pixel are then used to extract subregions of 9 × 9 pixels from the raw image. The size of subregion is set according to the PSF of our optical system.

#### Localization

We apply a maximum likelihood estimator to the W pixels in the subregion to calculate emitter location. From similar procedures in the literature [19], the parameters of a molecule 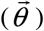 can be determined by minimizing the following equation:

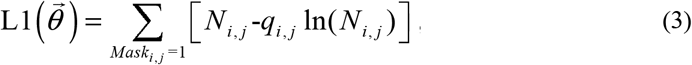

where N_i,j_ is the expected intensity, q_i,j_ is the observed intensity. In this study, N_i,j_ is set to be Gaussian distribution, where 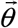 denotes the parameters that defines the Gaussian distribution, including amplitude (A), emitter location(x, y), and the standard deviation (s) which is determined by the point spread function (PSF) the optical system.

### 2.4 Determining emitter color for SRLM

For the colorimetry camera used in this study, the color of an emitter is encoded by four color channels (R, G, B, and NIR). Here we define normalized color intensity (NCI) to characterize the color of emitters detected in our imaging system. Note NCI is determined mainly by the emission spectrum of an emitter and the relative quantum efficiency curves of the camera (see Fig. 1(c)). NCI is calculated by the filtered intensity (color channels) divided by the unfiltered intensity (W channel). We calculate four NCI components (NCI_R, NCI_G, NCI _B, and NCI _NIR) to describe emitter color. Since different types of emitters have different emission spectra, the NCI components of these emitters should follow different distributions, which can be used for color recognition.

Singe-color SRLM experiments should be performed with all involved emitter types to build their corresponding NCI distributions. Later in multi-color SRLM analysis, the emitter color can be recognized and assigned by comparing the calculated NCI components of the emitters in subregions with the NCI distributions calculated from single-color SRLM experiments.

#### Calculating NCI components

From the Gaussian distribution determined in the localization step, we are able to recover the expected intensity (unfiltered) in color pixels. Ideally, after extracting the observed intensity (filtered) in these color pixels, we can acquire NCI components. However, after considering the effect of noise, we design a maximum likelihood estimator to calculate the NCI components.

Here we use r to represent the estimated NCI component. In a subregion, we assume there are K pixels in a color channel. For color pixel k (k∈[1,K]), the expected unfiltered intensity (recovered) is Z_k_, the expected filtered intensity is rZ_k_, and the observed filtered intensity is q_k_. Consequently, q_k_ follows Poisson distribution with a mean of rZ_k_. The joint probability function of this color channel in the subregion follows:

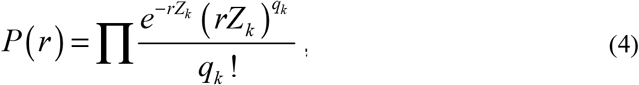

We can determine r with the maximum P by minimizing the following L function:

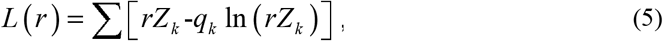

#### Building NCI distributions

By calculating the four NCI components from the same type of emitters, we are able to build the NCI distributions for different color channels. For each of the involved emitter types, we conduct single-color SRLM imaging, and build the corresponding NCI distributions.

#### Assigning emitter color

Considering that the four NCI components of an emitter are independent from each other (r_u_ is the estimated NCI component of the uth color channel, where u∈[1,U], U denotes the number of NCI components), for emitter type o (o∈[1,O]), the probability of one emitter arisen from a certain emitter type could be written as the product of the possibilities (normalized NCI distributions) of different channels:

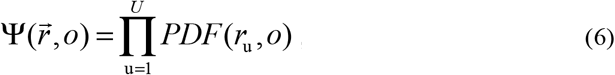

By normalizing and maximizing the probability above, we are able to determine the emitter color in a subregion.

### 2.5 Theoretical prediction of localization precision

Following the procedures by Thompson et al [20], here we derive an equation for predicting localization precision from the colorimetry camera. Although the W pixels for encoding emitter location cover only 50% of the pixels in a subregion, the intensity of a W pixel situated on position (i, j) still follows Gaussian distribution:

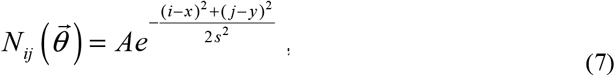

Where 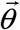 is a group of parameters that defines the Gaussian distribution, including amplitude (A), emitter location(x, y), and the standard deviation (s) which is determined by the point spread function (PSF) the optical system.

Estimating emitter location equals to minimizing the following sum of squared errors:

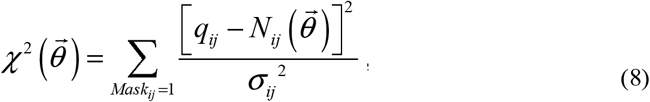

where q_ij_ is the observed emission intensity in pixel (i, j), N_ij_ is the expected emission intensity in pixel (i, j), and σ_ij_ is the uncertainty of N_ij_. Using Taylor expansion, we can write the mean squared error of θ_m_ as:

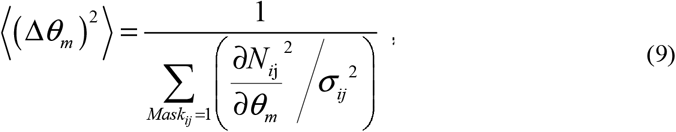

For the uncertainty brought by shot noise from the emitter:

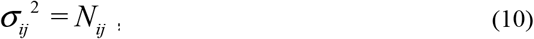

Hence, we obtain the localization error in one dimension (Δx) from shot noise by rewriting Eq. 9:

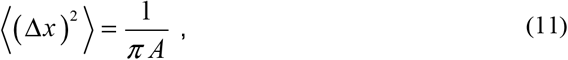

For the uncertainty induced by background, 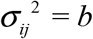, where b is the mean intensity of background, we calculate Δx from background as:

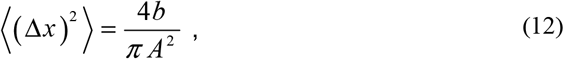

After considering detected signal, background, and pixelation noise, the localization precision in one dimension can be written as:

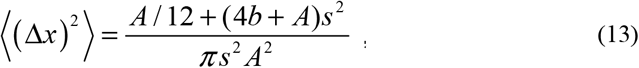

Using similar procedures, we can write the precision of standard deviation (Δs) and amplitude (ΔA) as:

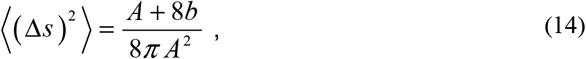

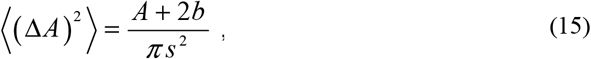

### 2.6 Theoretifcal prediction of NCI estimation error

Here we predict NCI estimation error (Δr) using the recovered unfiltered intensity (Z_k_ + b), the expected filtered intensity (rZ_k_+r_1_b), and the observed filtered intensity (q_k_). Calculating NCI estimation error equals to minimizing the following sum of squared errors [20]:

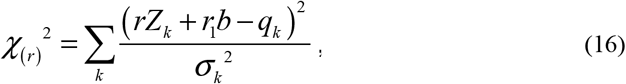

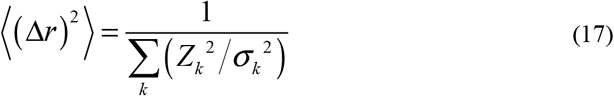

The intensity uncertainty of pixel k is from shot noise, localization error, and read noise:

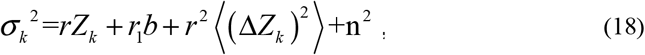

where n is the standard deviation of read noise, ΔZ_k_ is the intensity fluctuation brought by localization error.

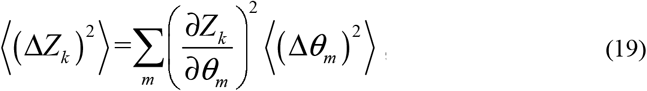

Substituting Eqs. 13-15 into Eq. 17, we can write NCI estimation error (Δr) as:

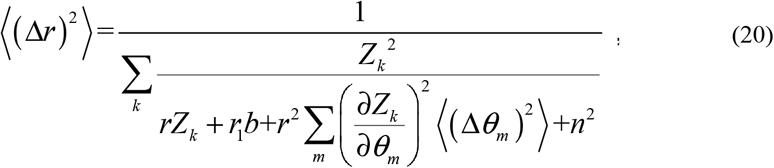

### 2.7 The procedures for determining emitter location and color

We employed SRLM as a representative application, and developed analysis procedures (Fig. 2) to decode emitter location and color from raw images. Taking a subregion containing only one emitter as an example, we firstly extract the W pixels from this subregion, and then use maximum likelihood estimator to calculate the emitter location. We further extract the color pixels from the same subregion, and estimate the normalized color intensity (NCI) of the emitter using the color pixels and the recovered Gaussian distribution from the emitter (see Section 2.3-2.4). Using the NCI values in four channels, we calculate the probability for identifying the color of the emitter. After repeating these decoding procedures (Fig. 2) in all acquired raw images, we obtain the locations and colors of a large number of emitters, and finally use them to reconstruct a two-color super-resolution (SR) image.

**Fig. 2.**
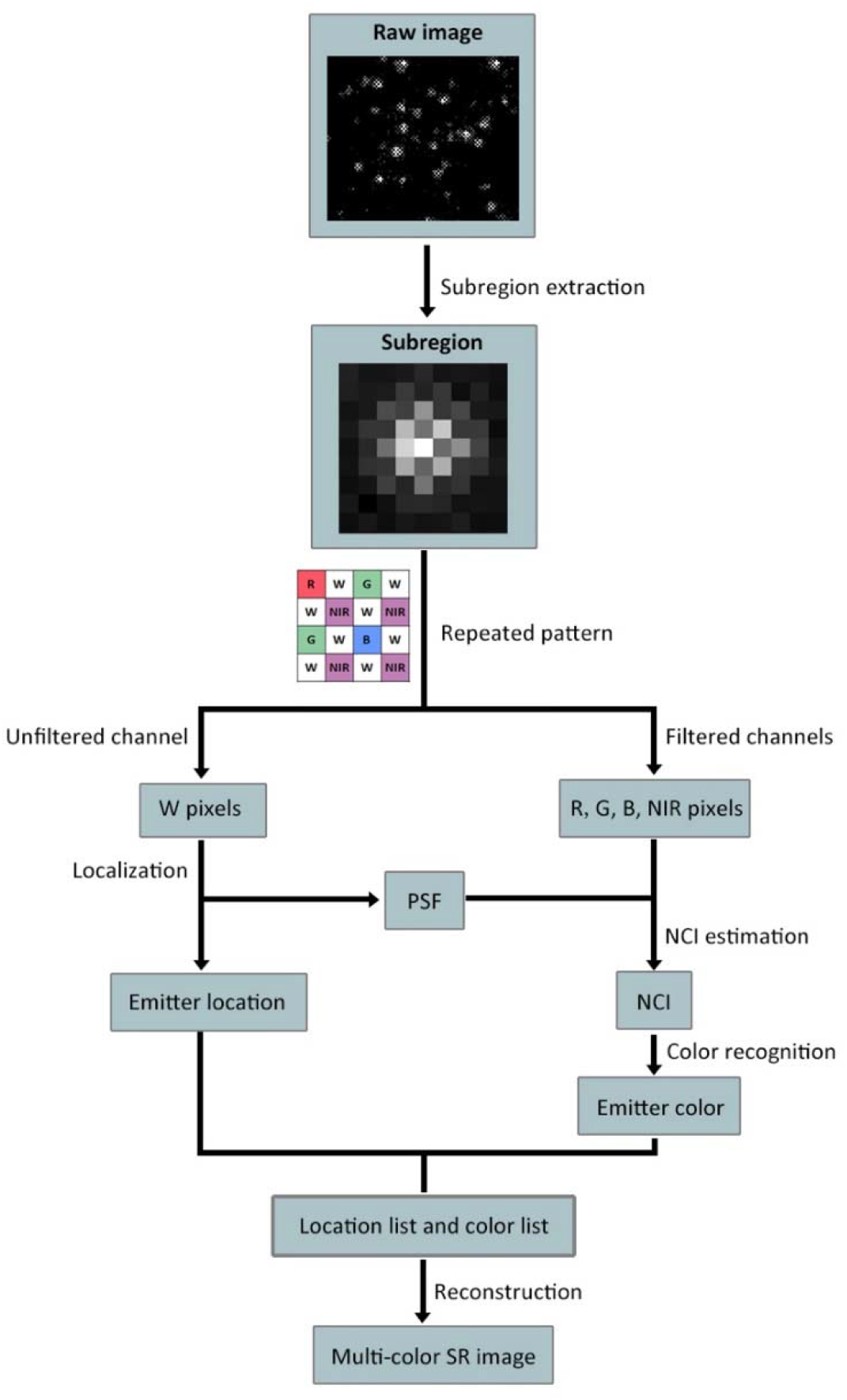
Data analysis procedures for super-resolution localization microscopy. This scheme displays only the procedures for one emitter in one raw image. These procedures should be repeated for all raw images to obtain a full list of emitter location and color, which is used to reconstruct a final multi-color super-resolution image.

## 3. Results

### 3.1 Determining emitter localization and color using simulated images

To quantify the performance of the colorimetry camera (Retina 200DSC, Tucsen Photonics) in estimating emitter location and color, we simulated raw images with a size of 64 × 64 pixels. For simplicity, we placed only one emitter in a single raw image. We quantified NCI estimation error and localization precision from a series of 1, 000 images in the following conditions: emission wavelength (400 - 700 nm), signal level (4,000 - 20,000 photon/molecule, typical for DL633 and CF680 in our experimental conditions due to the long exposure time), and readout noise (2.71 e-rms, measured for the colorimetry camera). We controlled the background level (336.4 photon/pixel), PSF standard deviation (1.34 pixel), and pixel size (108.3 nm) to match our SRLM experimental conditions. Because emitters can be centered on 16 different positions (see Fig. 1(b)), we also quantified NCI estimation error on these positions. We compared the localization precision of the colorimetry camera with Hamamatsu Flash 4.0 V3 (a popular monochrome camera used in SRLM). For the Flash 4.0 V3, readout noise is 1.6 e-rms, and QE is 0.78 in 660 nm.

We used simulated dataset to evaluate the estimation errors in emitter location and NCI. Note that 50% of the pixels in the colorimetry camera are W pixels, and the rest 50% are color pixels. Since the W pixels in a monochrome camera takes up all the pixels, the colorimetry camera suffers from a certain level of degradation in localization precision. We compared the localization precision of the colorimetry camera with a popular commercial low light camera (Hamamatsu Flash 4.0 V3), and found that the localization precision of the colorimetry camera (Eq. 13) is about 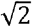 times that of the Flash 4.0 V3 (Fig. 3(a)). For the far-red emitters (DL633, CF680) used in this study, simulation shows that the localization precision is 4 nm for the Flash 4.0 V3, and 6 nm for the colorimetry camera. Furthermore, we confirmed that the central positions of the emitter in different kind of color pixels have negligible effect on the localization precision (Fig. 3(b)). We also investigated the relationship between the NCI estimation error and various photophysical parameters (signal level, position) (Fig. 3(c-d)). We found that the NCI estimation error basically follows the theoretical model (Section 2.6, Eq. 20), and that the central position of the emitter have negligible effect on NCI estimation.

**Fig. 3.**
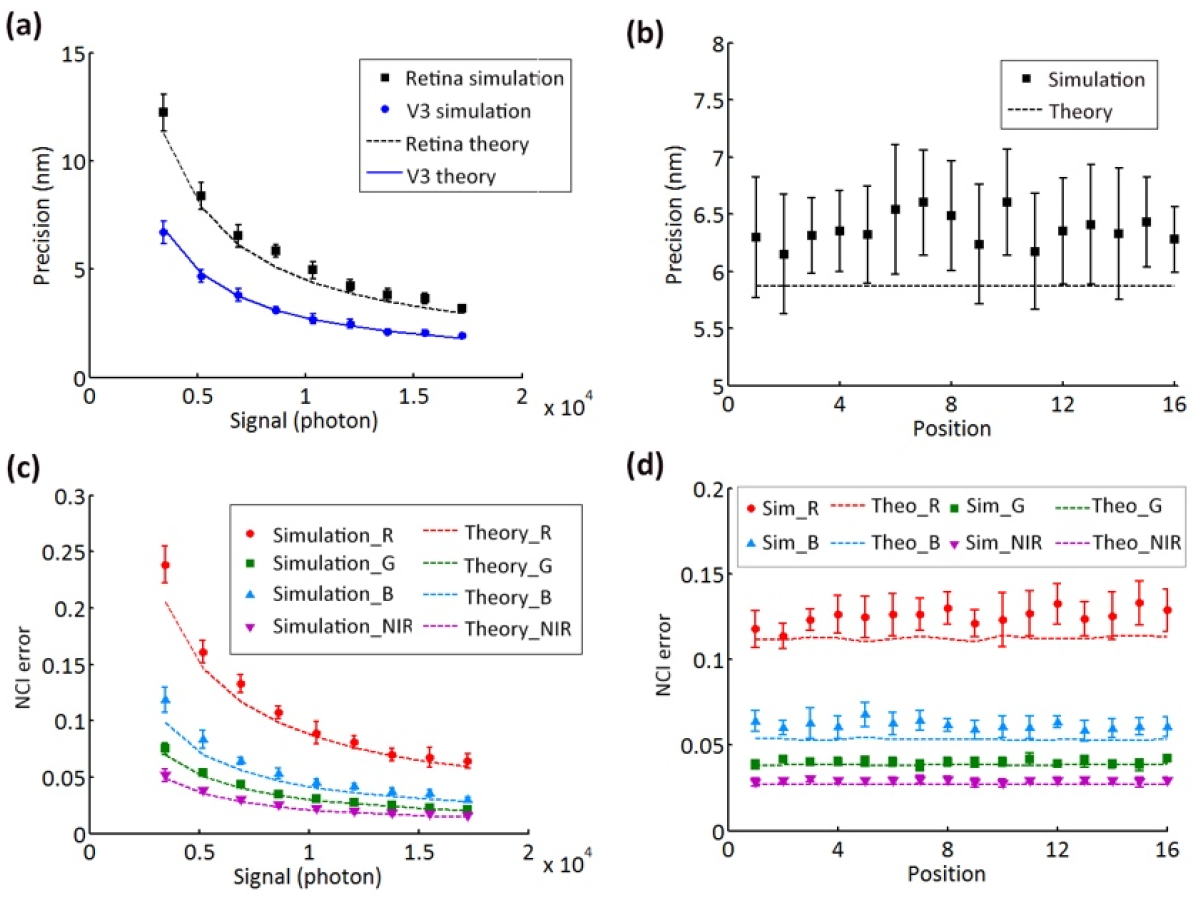
Performance of determining location and color on simulated dataset. (a) Localization precision under different signal intensities. (b) Localization precision for emitter situated on different pixel positions. (c) NCI estimation error under different signal intensities. (d) NCI estimation error for emitter situated on different pixel positions. Note that there are 16 different positions for a 4×4 pattern in the colorimetry camera, and that the positions shown in (b) and (d) point out the center locations of emitters. Error bars were from ten repeated measurements. The wavelength was 660 nm. Results for other wavelengths are similar.

### 3.2 Single-color SRLM

We verified the performance of our method using single-color SRLM on fixed cos-7 cells. Two representative SR images and the corresponding enlarged images are shown in Fig. 4(a-d), where microtubules were immunostained with DL633 (Fig. 4(a-b)) or mitochondria with CF680 (Fig. 4(c-d)). From a cross-sectional profile analysis, we obtained a maximum FWHM (full width at half maximum) resolution of 53.5 nm for the microtubule (see Fig. 4(e), the smaller value in the left), which is similar as the reported results [5]. Similar to the previously reported methods [4, 5], we used point-like objects in a SR image to calculate the image resolution inside cell sample. The localization precision from the colorimetry camera was experimentally measured to be 9-12 nm for these two emitters (see Fig. 5(a-c) for DL633, and Fig. 5(d-f) for CF680), which is sufficient to support SR imaging with ∼20 nm resolution.

**Fig. 4.**
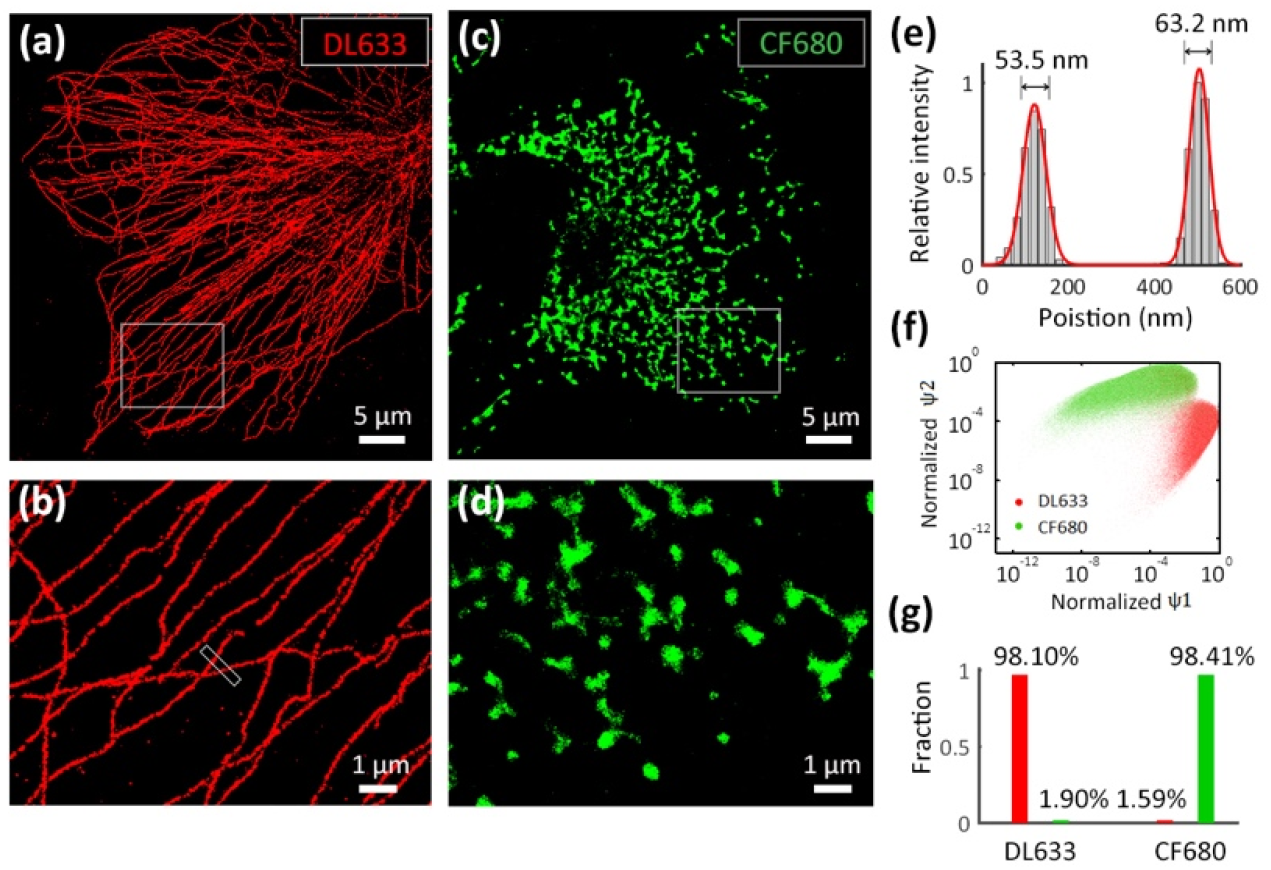
Single-color SRLM imaging. (a) SR image of microtubules in a fixed cos-7 cell labeled with DL633. (b) Zoom-in images of the rectangular areas in (a). (c) SR image of mitochondria in a fixed cos-7 cell labeled with CF680. (d) Zoom-in images of the rectangular areas in (c). (e) Cross-sectional profile of the boxed area in (b). The histograms (gray bars) were fitted with two Gaussian functions (red line). The FWHM resolution is shown on the top of the histograms. (f) Scatter plot of the normalized probability of an emitter arisen from a certain emitter type in SRLM. Here logarithmic scale is used. The plot is from 10, 000 experimental localizations detected in single color (DL633 or CF680) raw images. (g) Crosstalk between DL633 and CF680 (< 2%). Red bars represent the proportion of emitters that were identified as DL633. Green bars represent the proportion of emitters that were identified as CF680.

**Fig. 5.**
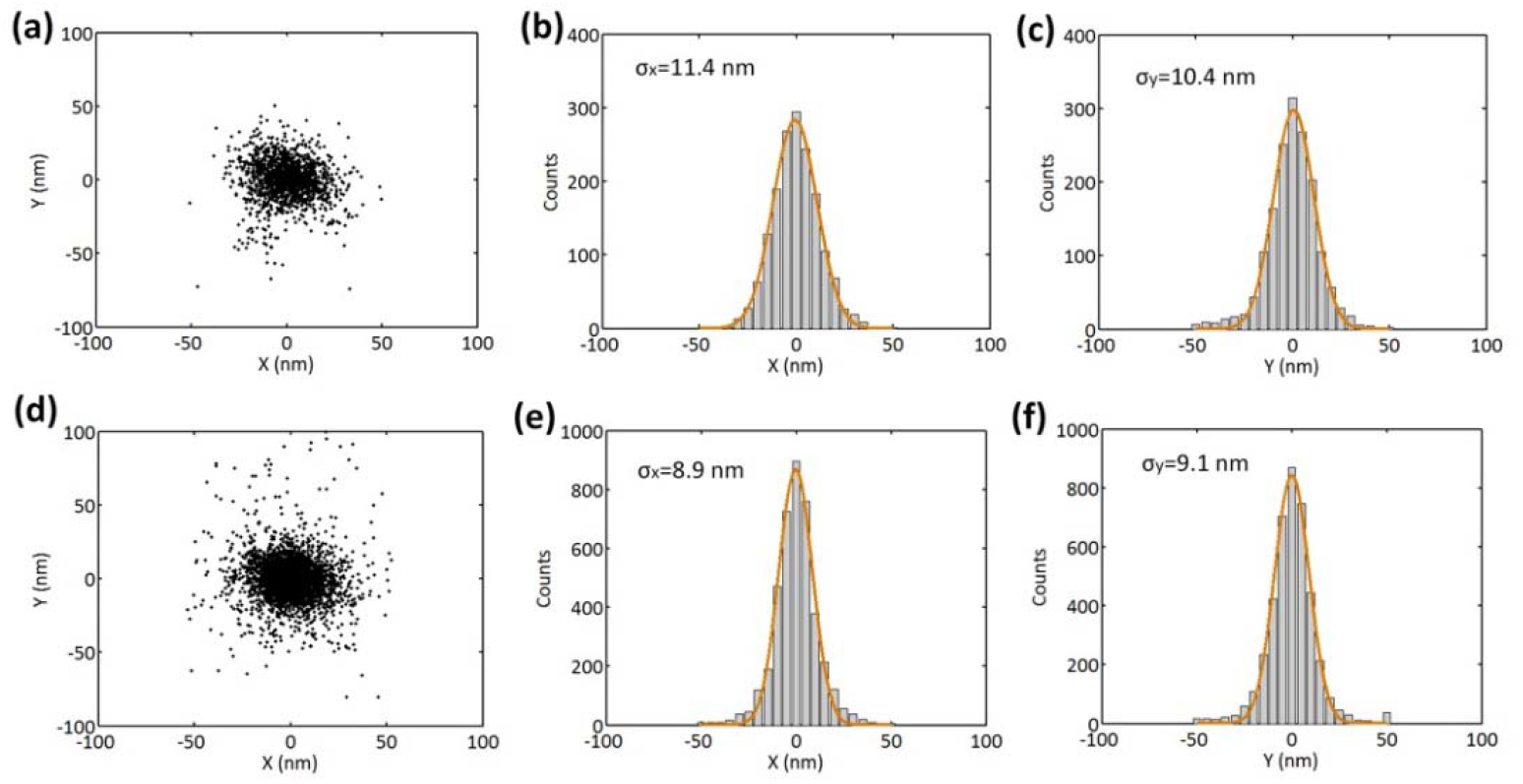
Localization distribution of point-like objects in DL633 and CF680 dataset. (a) Localization distribution of point-like objects in DL633 dataset. (b) The histogram of x dimension in (a). (c) The histogram of y dimension in (a). (d) Localization distribution of point-like objects in CF680 dataset. (e) The histogram of x dimension in (d). (f) The histogram of y dimension in (d). Clusters with > 9 emitters were aligned according to their center positions. The data were the same as those in Fig. 4. The distributions in (a) were from 1, 651 emitters in 99 clusters. The distributions in (d) were from 4, 136 localizations in 253 clusters. The FWHM resolution values are 26.8 nm (b), 24.5 nm (c), 21.0 nm (e) and 21.4 nm (f), respectively.

Given the fact that the cross-correlation drift correction method used in this study has a precision of 5-10 nm [21], the experimental precision of 9-12 nm is consistent with the theoretical precision of 6 nm in Section 3.1. We obtained a FWHM resolution of 21.0 nm (see Fig. 5(e), FWHM = 2.35σ) using the distribution of 4,136 localizations in 253 point-like objects. This FWHM resolution is close to the previously reported results [4, 5]. We assigned the color of each emitter to either DL633 or CF680 using the normalized probability distribution shown in Fig. 4(f). We also evaluated the misidentification between DL633 and CF680, and found that the crosstalk between DL633 and CF680 was < 2% (Fig. 4(g)), comparable to the lowest reported values in the literatures [4, 8].

### 3.3 Two-color SRLM

We carried out two-color SRLM on fixed cos-7 cells labeled simultaneously with DL633 (microtubules) and CF680 (mitochondria). We show a representative SR image and two enlarged images in Fig. 6(a-c). We analyzed the cross-sectional profiles of two microtubules and found an FWHM resolution of 54.7 nm and 60.6 nm (Fig. 6(d-e)), respectively. Additionally, in our method, the minimum distance between emitters is found to be 9 pixels, which is the subregion size covered by one emitter. Therefore, for the reported multi-color methods with low crosstalk (< 2%), the minimum separation distance in our method is the same as that in salvaged fluorescence based method [8], but is significantly shorter than that in the chromatic dispersion based methods (∼20 pixel) [4, 9]. This finding indicates that our method allows high emitter density in raw images, thus enabling good potential in high-throughput multi-color SRLM.

**Fig. 6.**
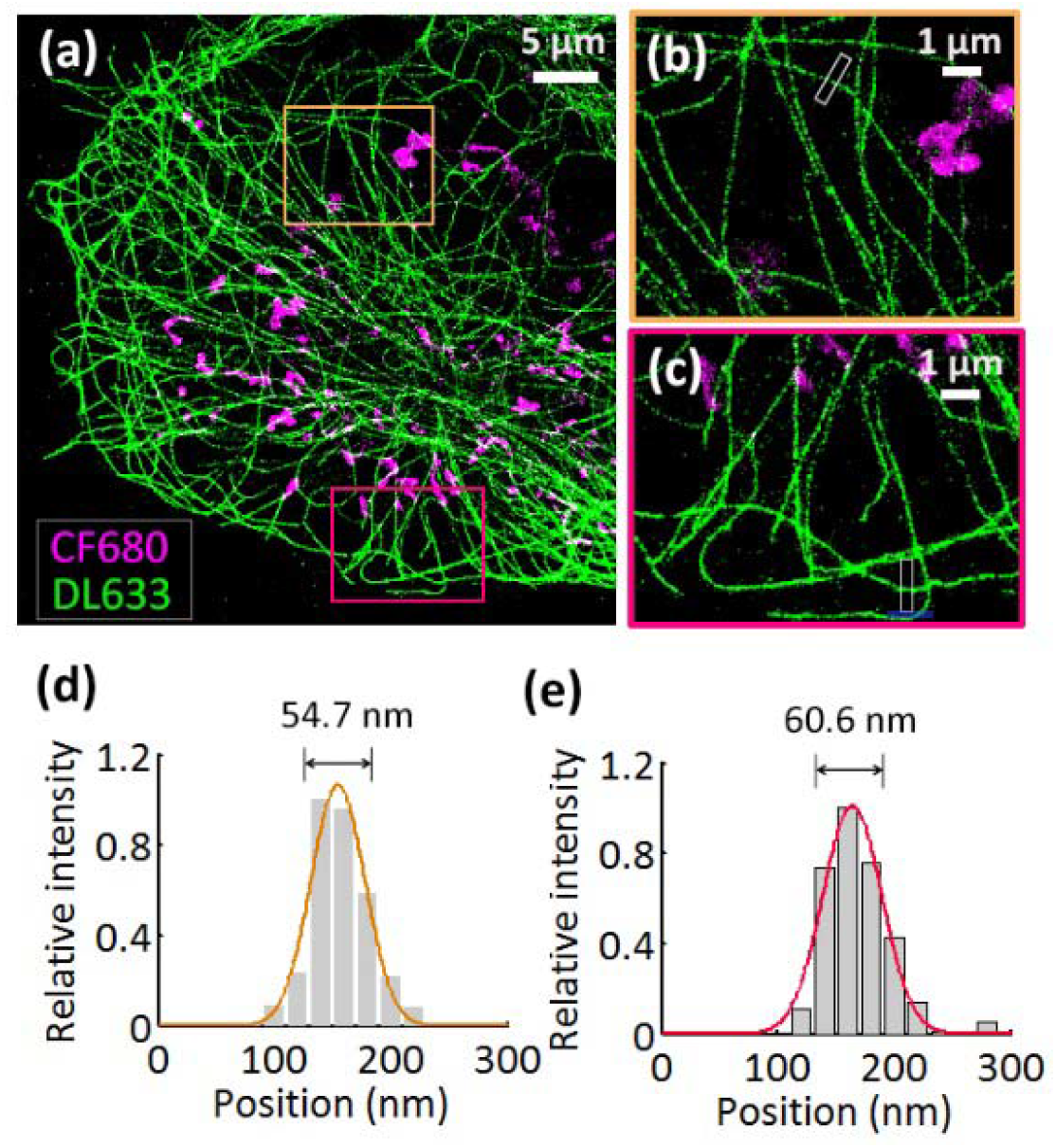
Simultaneous two-color SRLM. (a) A reconstructed SR image of a fixed cos-7 cell labeled with DL633 (microtubules) and CF680 (mitochondria), respectively. (b-c) Zoom-in images of the rectangular areas in (a). (d-e) Cross-sectional profiles of the boxed areas in (b-c). The histograms (gray bars) were fitted with Gaussian function (orange and red lines). The FWHM resolution is shown on the top of the histograms.

## 4. Discussion

We proposed a new method for two-color super-resolution localization microscopy. By taking advantage of the repeated pixel pattern of a customized RGBW camera (called colorimetry camera in this study), we are able to realize joint encoding of emitter location and color in a single raw image. We demonstrated that two-color SRLM is possible by combining a colorimetry camera and a basic optical set-up. We further verified that our method is capable of providing both low crosstalk (< 2%) and high image resolution (∼20 nm), which are comparable to the best reported values.

The major disadvantages of our method is from two sources: (1) the current colorimetry camera has a moderate quantum efficiency (< 70% for the white pixels and <60% for the color pixels), limiting the image quality and thus the localization precision and the spectral discrimination ability; (2) only the white pixels (which occupy 50% of all pixels) are used for molecule localization, resulting in under-sampling and decreased localization precision. The limitation in quantum efficiency may be partially overcome by developing back-illuminated colorimetry cameras. And the localization precision issue can be minimized by developing new algorithms which use all pixels for molecule localization.

It is worthy to note that here we only demonstrated 2D SRLM with the colorimetry camera; however, in principle, our method can be expanded to 3D SRLM if we make appropriate modification in optical set-up (for example, including a cylindrical lens in the detection path). And, with the selection of appropriate emitters, our method may be able to realize simultaneous SRLM with 3∼6 colors. Due to the limited spectral discrimination ability of our method (∼40 nm), a possible way to achieve 3-6 color SRLM is to screen emitters which have (1) well-separated emission; (2) good photophysical properties for SRLM; (3) low fluorescence background from simultaneous excitation of multiple lasers.

In summary, we demonstrated simultaneous two-color SRLM imaging using two common emitters (DL633 and CF680, with 40 nm separation in emission maximum) and achieved a low crosstalk of < 2%. In comparison, the PSF engineering based method, which is the only reported method for simultaneous multi-color SRLM, was able to discriminates two emitters with ∼100 nm emission separation and ∼20% crosstalk. Compared with the chromatic dispersion based methods (which are capable of distinguishing four emitters with ∼10 nm emission separation and < 2% crosstalk, but require an elongated PSF), our method is poorer in the spectral discrimination ability, but has the desirable performance of low crosstalk, and allows a larger activation density (via no elongated PSF). After considering the simplicity of our method (uses a basic optical set-up and requires no image registration), we believe this study will encourage a popular use of multi-color SRLM in biomedical researches, and stimulate a rising interest in the development of new sCMOS cameras for multi-color imaging at low light condition.

## Acknowledgments

We thank the Optical Bioimaging Core Facility of WNLO-HUST for technical support, and Tucsen Photonics for customizing the sCMOS camera.

## Data availability

Data underlying the results presented in this paper are not publicly available at this time but may be obtained from the authors upon reasonable request.

## References

1. Y. M. Sigal, R. Zhou, and X. Zhuang, “Visualizing and discovering cellular structures with super-resolution microscopy,” Science 361, 880–887 (2018).

2. S. J. Sahl, S. W. Hell, and S. Jakobs, “Fluorescence nanoscopy in cell biology,” Nat. Rev. Mol. Cell Biol. 18, 685–701 (2017).

3. P. A. Gómez-García, E. T. Garbacik, J. J. Otterstrom, M. F. Garcia-Parajo, and M. Lakadamyali, “Excitation-multiplexed multicolor superresolution imaging with fm-STORM and fm-DNA-PAINT,” Proc. Natl Acad. Sci. USA 115, 12991–12996 (2018).

4. Z. Zhang, S. J. Kenny, M. Hauser, W. Li, and K. Xu, “Ultrahigh-throughput single-molecule spectroscopy and spectrally resolved super-resolution microscopy,” Nat. Methods 12, 935–938 (2015).

5. M. Bates, B. Huang, G. T. Dempsey, and X. Zhuang, “Multicolor super-resolution imaging with photo-switchable fluorescent probes. Science 317, 1749–1753 (2007).

6. G. T. Dempsey, J. C. Vaughan, K. H. Chen, M. Bates, and X. Zhuang, “Evaluation of fluorophores for optimal performance in localization-based superresolution imaging,” Nat. Methods 8, 1027–1036 (2011).

7. I. Testa, C. A. Wurm, R. Medda, E. Rothermel, C. Von Middendorf, J. Fölling, S. Jakobs, A. Schönle, S. W. Hell, and C. Eggeling, “Multicolor fluorescence nanoscopy in fixed and living cells by exciting conventional fluorophores with a single wavelength,” Biophys. J. 99, 2686–2694 (2010).

8. Y. Zhang, L. K. Schroeder, M. D. Lessard, P. Kidd, J. Chung, Y. Song, L. Benedetti, Y. Li, J. Ries, J. B. Grimm, L. D. Lavis, P. D. Camilli, J. E. Rothman, D. Baddeley, and J. Bewersdorf, “Nanoscale subcellular architecture revealed by multicolor three-dimensional salvaged fluorescence imaging,” Nat. Methods 17, 225–231 (2020).

9. B. Dong, L. Almassalha, B. E. Urban, T. Q. Nguyen, S. Khuon, T. L. Chew, V. Backman, C. Sun, and H. F. Zhang, “Super-resolution spectroscopic microscopy via photon localization,” Nat. Commun. 7, 12290 (2016).

10. Y. Shechtman, L. E. Weiss, A. S. Backer, M. Y. Lee, and W. E. Moerner, “Multicolour localization microscopy by point-spread-function engineering,” Nat. Photonics 10, 590−594 (2016).

11. E. Hershko, L. E. Weiss, T. Michaeli, and Y. Shechtman, “Multicolor localization microscopy and pointspread-function engineering by deep learning,” Opt. Express 27, 6158–6183 (2019).

12. K. S. Grußmayer, S. Geissbuehler, A. Descloux, T. Lukes, M. Leutenegger, A. Radenovic, and T. Lasser, “Spectral cross-cumulants for multicolor super-resolved SOFI imaging,” Nat. Commun. 11, 3023 (2020).

13. A. von Diezmann, Y. Shechtman, and W. E. Moerner, “Three-dimensional localization of single molecules for super-resolution imaging and single-particle tracking,” Chem. Rev. 117, 7244–7275 (2017).

14. T. Rahbek-Clemmensen, M. D. Lycas, S. Erlendsson, J. Eriksen, M. Apuschkin, F. Vilhardt, T. N. Jørgensen, F. H. Hansen, and U. Gether, ”Super-resolution microscopy reveals functional organization of dopamine transporters into cholesterol and neuronal activity-dependent nanodomains,” Nat. Commun. 8, 740 (2017).

15. Y. Zhang, M. Lara-Tejero, J. Bewersdorf, and J. E. Galán, “Visualization and characterization of individual type III protein secretion machines in live bacteria,” Proc. Natl Acad. Sci. USA 114, 6098–6103 (2017).

16. R. D. Jansen□van Vuuren, A. Armin, A. K. Pandey, P. L. Burn, and P. Meredith, “Organic Photodiodes: The Future of Full Color Detection and Image Sensing,” Adv. Mater. 28, 4766–4802 (2016).

17. W. Choi, H. Park, and C. Kyung, “Color reproduction pipeline for an RGBW color filter array sensor,” Opt. Express 28, 15678–15690 (2020).

18. S. Van De Linde, A. Löschberger, T. Klein, M. Heidbreder, S. Wolter, M. Heilemann, and M. Sauer, “Direct stochastic optical reconstruction microscopy with standard fluorescent probes,” Nat. Protoc. 6, 991–1009 (2011).

19. T. W. Quan, P. Li, F. Long, S. Zeng, Q. Luo, P. N. Hedde, G. U. Nienhaus, and Z. L. Huang, “Ultra-fast, high-precision image analysis for localization-based super resolution microscopy,” Opt. Express 18, 11867–11876 (2010).

20. R. E. Thompson, D. R. Larson, and W. W. Webb, “Precise Nanometer Localization Analysis for Individual Fluorescent Probes,” Biophys. J. 82, 2775–2783 (2002).

21. Y. Wang, J. Schnitzbauer, Z. Hu, X. Li, Y. Cheng, Z. L. Huang, and B. Huang, “Localization events-based sample drift correction for localization microscopy with redundant cross-correlation algorithm,” Opt. Express 22, 15982–15991 (2014).

